# A Comprehensive and Accessible Model for Co-Segregation Analysis in *BRCA1, BRCA2*, and *PALB2* Variant Classification

**DOI:** 10.1101/2024.09.23.614309

**Authors:** Setareh Moghadasi, Ramin Monajemi, Merel E. Braspenning, Maaike P.G. Vreeswijk, Mar Rodríguez-Girondo

## Abstract

Variants of uncertain significance (VUS) are genetic variations with unclear clinical implications, often complicating clinical management in genetic testing. The analysis of co-segregation of the variant with the disease in families has been shown to be a powerful tool for the classification of these variants. We present CAL-Leiden (Co-segregation Analysis via Likelihood ratio analysis-Leiden), a comprehensive co-segregation model facilitating the classification of variants in *BRCA1, BRCA2* and *PALB2* genes, which can be used as an important component of the ACMG/AMP classification guideline. CAL-Leiden includes an expanded range of cancer types, including pancreatic cancer, in addition to breast and ovarian cancer. It also considers contralateral breast cancer. The model integrates population incidence rates from the Netherlands and the United Kingdom, along with penetrance data from the latest literature. A web-based platform has been developed, making the model accessible and practical for use in diagnostic labs: https://bioexp.net/cosegregation/. We demonstrate the functionality of the tool with multiple pedigrees and compare its performance with alternative approaches. These features in CAL-Leiden collectively contribute to a more comprehensive and accurate assessment of variant pathogenicity, helping lab specialists in classification of the variants of uncertain significance.

## INTRODUCTION

The broad application of sequencing technologies in DNA diagnostic laboratories and the use of extensive cancer related gene panels have resulted in the identification of a growing number of sequence variants in major cancer-predisposition genes such as *BRCA1, BRCA2* and *PALB2*, of which the clinical importance remains uncertain, referred to as Variants of Uncertain Significance (VUS). A variety of approaches have been used to assess the clinical relevance of these VUS, among which co-segregation analysis.

Co-segregation analysis studies the inheritance of a genetic variant within a family, particularly its presence in affected or unaffected family members, to determine the likelihood of its pathogenicity or association with a specific disease. Different statistical approaches have been described to determine the likelihood of pathogenicity using co-segregation data which can then be integrated together with supplementary evidence for the application in ACMG/AMP-based classification in which all the available data against or in favor of pathogenicity are combined (1).

The major contribution in the early 2000s was the introduction of the full pedigree likelihood model (2). This method proposed the concept of assessing the probability that observed genetic patterns within family pedigrees align with a predefined gene-disease penetrance model. The method employed incidence rates to depict the penetrance for affected individuals. However, these might not be entirely suitable for age-dependent phenotypes such as cancer age of onset. Moreover, within the full pedigree approach, the penetrance was assumed to remain constant across 10-year time intervals (liability classes), which may not accurately reflect actual biology. Additionally, its implementation relied on specialized linkage software, which was not practical for the general user. In 2009, Mohammadi et al. (3) in Leiden, the Netherlands, introduced a novel pedigree likelihood model based on a survival analysis approach. In contrast to the full pedigree likelihood model, which examines data solely at the time of diagnosis, the survival approach considers the individual’s entire lifespan. This implies that affected individuals contribute to the model through a density function rather than an incidence rate and unaffected individuals contribute with the survival function at last follow-up. The density function, capturing both disease-free intervals and cancer diagnoses, contains richer information, offering a more comprehensive perspective on genetic penetrance. Additionally, the Mohammadi model simultaneously considered breast cancer (BC), ovarian cancer (OC), and contralateral breast cancer (CBC) as contributing phenotypes to the penetrance model. Moreover, a user-friendly web-based tool was made available, facilitating the analysis of pedigrees by users not skilled with specialized software (3).

Recently, two papers were published using a similar survival approach. Rañola et al. (4) presented an adaptation of the Mohammadi model (3), integrating the latest age-dependent BC penetrance data for *BRCA1* and *BRCA2* carriers and other genes such as *ATM, CHEK2, MEN1, MLH1, MSH2, MSH6*, and *PMS2* were included. However, this adaptation simplified the original Mohammadi penetrance model; focusing solely on the first cancer, diminishing the original comprehensiveness of the Mohammadi’s penetrance model. More recently, Belman et al. (5) revised the original full pedigree likelihood model by incorporating a survival formulation in the likelihood specification resulting in a similar statistical formulation to the Mohammadi model (3).

This study introduces a new model based on Mohammadi et al. (3), incorporating several significant enhancements. These include the addition of *PALB2* alongside *BRCA1* and *BRCA2*, and the expansion of considered cancers beyond breast and ovarian to include pancreatic cancer. Moreover, updates to population incidence rates, using more recent Dutch and United Kingdom (UK) data, and adjustments to penetrance values based on the latest literature have been incorporated. A new website (https://bioexp.net/cosegregation/) has been launched. We evaluated the tool with multiple pedigrees and compared the results with alternative approaches by Rañola et al. (4) and Belman et al. (5). These enhancements contribute to a more reliable assessment of variant pathogenicity and help the lab specialists in the classification of these variants.

## SUBJECTS AND METHODS

### Mohammadi et al. (3) (2009) revised and extended

The likelihood ratio (LR) for pathogenicity of a variant based on segregation within families is defined as the probability of observing the family profile of variant genotypes given the disease phenotypes, assuming that the variant under investigation is pathogenic, divided by the probability of observing the family profile of variant genotypes given the disease phenotypes assuming that the variant under investigation is benign:

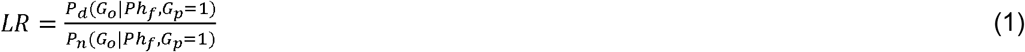

where *Ph*_*f*_ stands for the phenotypes in family *f, G*_*O*_ stands for the observed profile of genotypes of the VUS in family *f, and G*_*p*_ is the genotype of the proband, who is a carrier by definition. The overall objective is to elucidate whether the observed phenotypes in the family are supported by the hypothesis that the variant is benign or pathogenic. In case of benignity, one would expect *P*_*d*_(*G*_*o*_ | *Ph*_*f*_,*G*_*p*_ =1) to be similar to *P*_*n*_(*G*_*o*_ | *Ph*_*f*_,*G*_*p*_ =1) and the *LR* will be close to1. Alternatively, when pathogenic, one would expect that *P*_*d*_(*G*_*o*_ | *Ph*_*f*_,*G*_*p*_ =1) > *P*_*n*_(*G*_*o*_ | *Ph*_*f*_,*G*_*p*_ =1) and hence a *LR*>1. Increasing values of *LR* indicate increasing evidence of the compatibility of the observed family phenotypes and a pathogenic variant in the gene under investigation. The technical details can be found in Mohammadi et al. (3) and in supplementary material, appendix A. It is important to emphasize that the key components for the calculation of the *LR* are the likelihood contributions for each family member *i* under the assumptions of variant benignity and pathogenicity based on their phenotypical information. Once a specification for the individual likelihood contributions is available, the model allows to reconstruct the joint, family-specific, likelihood contributions used in expression (1) above.

The individual-specific phenotypical information is the age of onset of the cancers associated with the gene under investigation or the age at the last examination for unaffected individuals. In the Mohammadi model (3), the age of onset of BC, OC and CBC were considered. This meant that the basic phenotypical information for female *i* is given by (*t*_*bci*_,*t*_*ovi*_,*t*_*cbci*_,*d*_*bci*_,*d*_*ovi*,_*d*_*cbci*_), where *d*_*bci*_ is the BC affection status indicator, *d*_*bci*_=1 if female *i* is affected by BC and 0 otherwise and *t*_*bci*_ represents the age of BC diagnosis for the affected and the age of the last examination for unaffected individuals. Analogously, *t*_*ovi*_ represents the age at OC diagnosis for the affected and the age of the last examination for unaffected individuals; *d*_*ovi*_ is the OC affection status indicator, *d*_*ovi*_=1 if female *i* has been diagnosed with OC and 0 otherwise. Additionally, *t*_*cbci*_, where *t*_*cbci>*_*t*_*bci*,_ represents the age at CBC diagnosis and *d*_*cbci*_ its corresponding affection status indicator. If unaffected at the end of follow-up, *t*_*bci*_=*t*_*ovi*_ =*t*_*cbci*_ are the age at the last examination and *d*_*bci*_=*d*_*ovi*_=*d*_*cbc*i=_0. In absence of CBC, assuming independence of BC and OC diagnosis, and based on standard likelihood theory for survival data, the likelihood contribution of female *i* is given by:

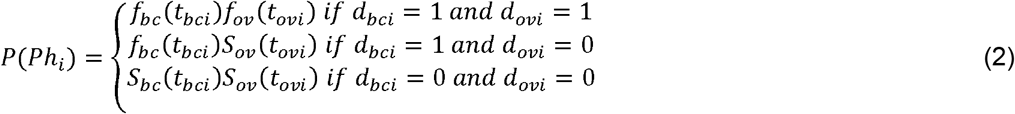

Where *S*_*bc*_(*t*_*bci*_)=1-*F(t*_*bci*_) and *S(t*_*ovi*_)=1-*F(t*_*bci*_) are the probabilities of being free of BC and OC by age *t*_*bci*_ and *t*_*ovi*._. *F*_*bc*_*(t*_*bci*_) and *F*_*ov*_*(t*_*ovi*_) are the penetrances, i.e. the cumulative risks of having BC and OC before ages *t*_*bci*_ *and t*_*ovi*_, and *f*_*bc*_*(t*_*bci*_) and *f*_*ov*_*(t*_*ovi*_) are their corresponding derivatives, the density functions. Specific values of these functions under the assumptions of pathogenicity and benignity are available from external sources and plugged-in (2) to obtain the two factors of expression (1). Note that for each of the cancers under investigation, the density function at time *t*_*i*_ can be rewritten as *f(t*_*i*_*)=h(t*_*i*_*)S(t*_*i*_), making evident the intuitive idea that the phenotypical information for an individual *i* affected with certain cancer type at age *t*_*i*_ is that *i* was free of that type of cancer until age *t*_*i*_ and that the event was observed at *t*_*i*_ (this instantaneous probability is given by *h(t*_*i*_), also known as the hazard and can be approximated by the incidence rate). The penetrance functions used by Mohammadi et al. (3) were smoothed adaptations of those described in the state-of-the art literature of the time, based on normal distribution functions (6). Contralateral breast cancer, when present, was considered by modifying the term *f*_*bc*_*(t*_*bci*_) in expression (2) and replacing it by 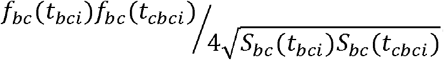. This modification assumed that the occurrence of cancer in each breast is an independent process. At that time, there was not available information on the conditional probability of CBC; thus, the available penetrance of BC was assumed to represent the minimum of these two independent processes (see supplementary material, appendix B for details). The independence assumption in expression (2) implies that after the diagnosis of one gene-related cancer, the risk of a second cancer of interest remains the same as it was before the first cancer diagnosis. This assumption is arguable since one can expect that the risk of a second cancer diagnosis might be modified by the first diagnosis due to changes in patient surveillance and treatment effects. Given the absence of reliable data on these conditional risks, one might choose to end the follow-up after the first diagnosis to avoid bias. Consequently, the age at onset of the remaining relevant cancers is right censored at the age of the first cancer diagnosis. The Mohammadi model (3) could also be applied in this scenario, provided the dataset was appropriately modified by the user.

Expression (2) aligns with the penetrance formulation proposed by Belman et al. (5), which uses the same phenotypic likelihood formulation but enforces stopping follow-up after the first cancer diagnosis. The approach implemented by Rañola et al. (4) is less comprehensive. In their model, either primary BC or OC, whichever occurs first, is used to define the affection status. However, the penetrance for unaffected individuals solely considers BC. Neither Belman (5), nor Rañola (4) models allow the inclusion of CBC. Derivations and a detailed model comparison among the three methods can be found in the supplementary material, appendix C.

### Extension of the Mohammadi model: more genes, cancer types, and updated penetrances

The Mohammadi model (3) has undergone expansion in three directions: first, by including the *PALB2* gene and second, by incorporating additional cancer types into the algorithm, third by updating the penetrances based on the latest literature. The integration of multiple cancers is explained for the case BC and OC in expression (2) above. Furthermore, age-specific cancer incidence rates and corresponding cumulative risks used in expression (2) have been used in accordance with the latest literature. Five-year age-specific cancer incidence rates for carriers of pathogenic variants in *BRCA1, BRCA2*, and *PALB2* were computed by multiplying the underlying population incidence rates by the most recently published age-specific relative risks (7-12). A comprehensive list of the considered cancers per gene, along with the updated parameters and references, can be found in supplementary material tables 1 and 2. We considered two possible reference populations: The Netherlands and the United Kingdom. We used data from IKNL for the year 2017 for the Netherlands (13), and for the UK, data corresponding to the period 2015-2017 were included (14). Similar to the initial proposal by Mohammadi et al. (3), we considered smoothed versions of the penetrance based on a normal distribution assumption for the cancer incidence data.

For CBC, we introduced a novel approach. Using available data on conditional relative risks of CBC given the age of primary of BC for *BRCA1* and *BRCA2* carriers (7) and in the general population (15), we developed a new method to model the penetrance of CBC based on the following expression, which does not assume independence:

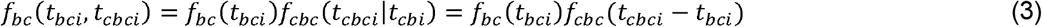

Similarly to other cancers, we applied a smoothing approach to the term *f*_*cbc*_ (*t*_*cbci*_ − *t*_*bci*_) based on an exponential, constant-hazards distribution, given that corresponding raw incidence rates do not clearly vary with age. Due to the limited number of events among carriers, penetrance values stratified by age groups of primary BC were not considered (7). Further details on the modeling of CBC can be found in supplementary material, appendix B.

### A new web application

A user-friendly website has been created, featuring pedigree visualization and intuitive data entry with a brief report generation functionality (https://bioexp.net/cosegregation/). The pedigree input format aligns with the data format used in CanRisk (https://www.canrisk.org/), a web-based tool widely employed for personalized risk assessments of BC and OC (16). CanRisk is increasingly used as a medical device in international clinical practices, and as a result, the input file for tested families is frequently available. To enhance user convenience, we have chosen to adopt the same data format for input on our website. An additional enhancement is that CAL-Leiden provides users with the option to consider all gene-related cancers within one person, account for CBC, or stop the person-specific follow-up after the first gene-related cancer diagnosis, all via a drop-down menu option without modifying the input data.

CAL-Leiden, as in the Mohammadi et al. (3), relies on the same computational algorithm that enumerates all genotypic configurations of a rare variant. Compared to the Mohammadi website application, this model demonstrates an improved computational performance and reduced memory consumption. However, in some large or complex pedigrees, CAL-Leiden may be unable to calculate the LR due to this algorithm’s inherent limitations. In such cases, we recommend trimming branches with unaffected and/or untested individuals, in line with Mohammadi et al. (3). Specifically, this includes family members younger than 20 years who are unaffected by relevant types of cancer or are untested. Privacy and security considerations were considered during the website application’s development. Pedigrees uploaded to the website are not retained. Users can generate a PDF-report of the analysis for their own records. Furthermore, the code is available upon request. For more information, please contact the corresponding author.

## RESULTS

After implementing the updates described in the methods section, we conducted a side-by-side comparison between the output of CAL-Leiden, Mohammadi et al. (3), COOL (version 2) (5), and the Rañola et al. (4) approach as implemented in the R package Coseg and website analyzemyvariant.org, respectively. For clarity and ease of comparison, our primary analysis was based on the same pedigree described in Belman et al. (5). We revisited the impact of different affection and carrier status of the 81-year-old female family member (Figure 1, Table 1) and extended the analysis by varying her age from 20 to 85 years (Figure 2).

**Figure 1.**
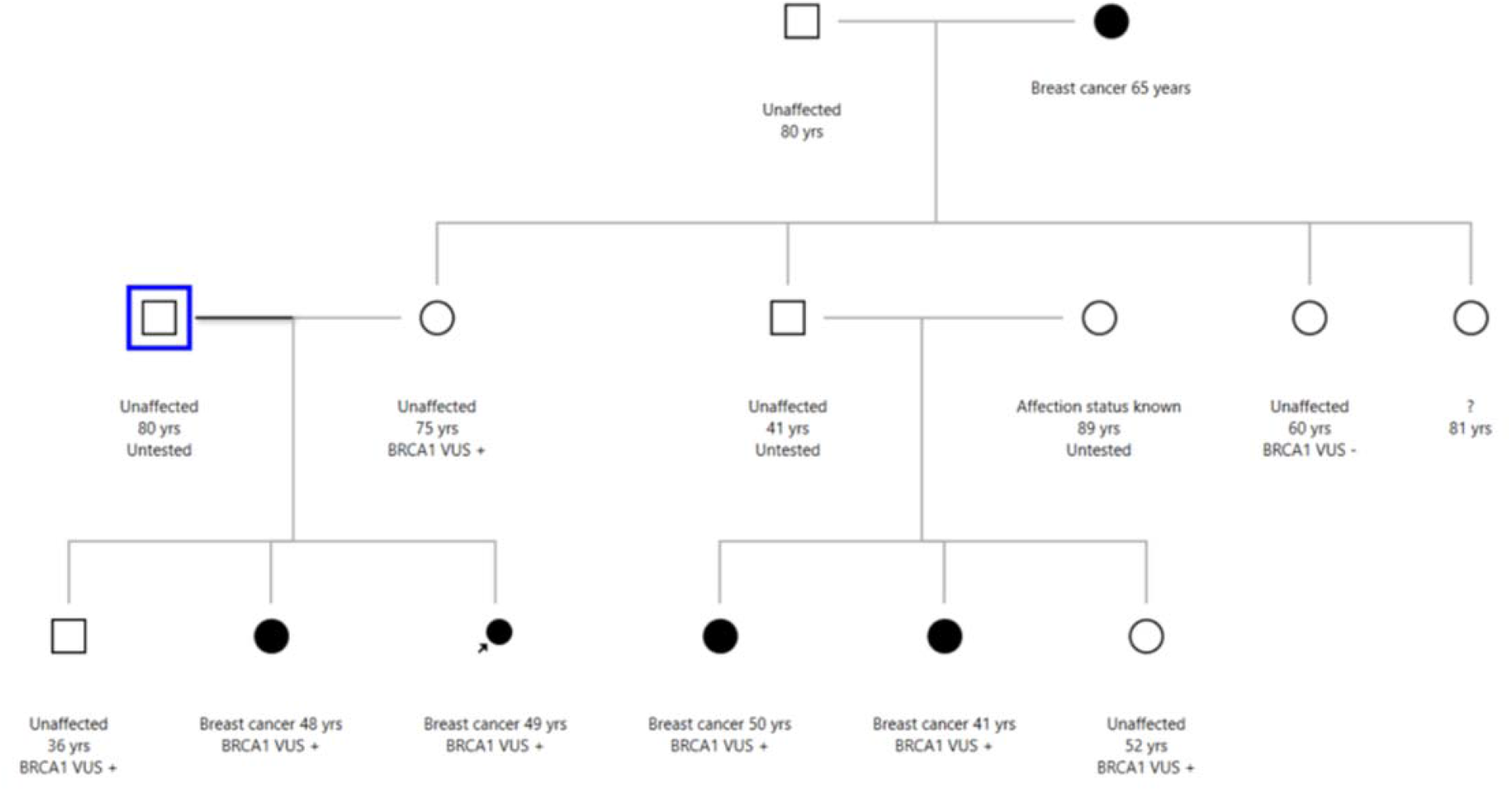
Figure 1 is based on figure 1 in Belman et al (5). In this pedigree Proband is indicated with an arrow Black filled: affected with breast cancer. Age of onset for affected and age of the last follow-up for unaffected individuals is indicated. Genotype is indicated as “BRCA1 VUS+ for carrier of the VUS BRCA1 VUS-for not carriers of the VUS in the BRCA1. For person named “?” affection status and genotype is to be determined. The default data in COOL and the United Kingdom population penetrance data in CAL-Leiden are used.

**Figure 2.**
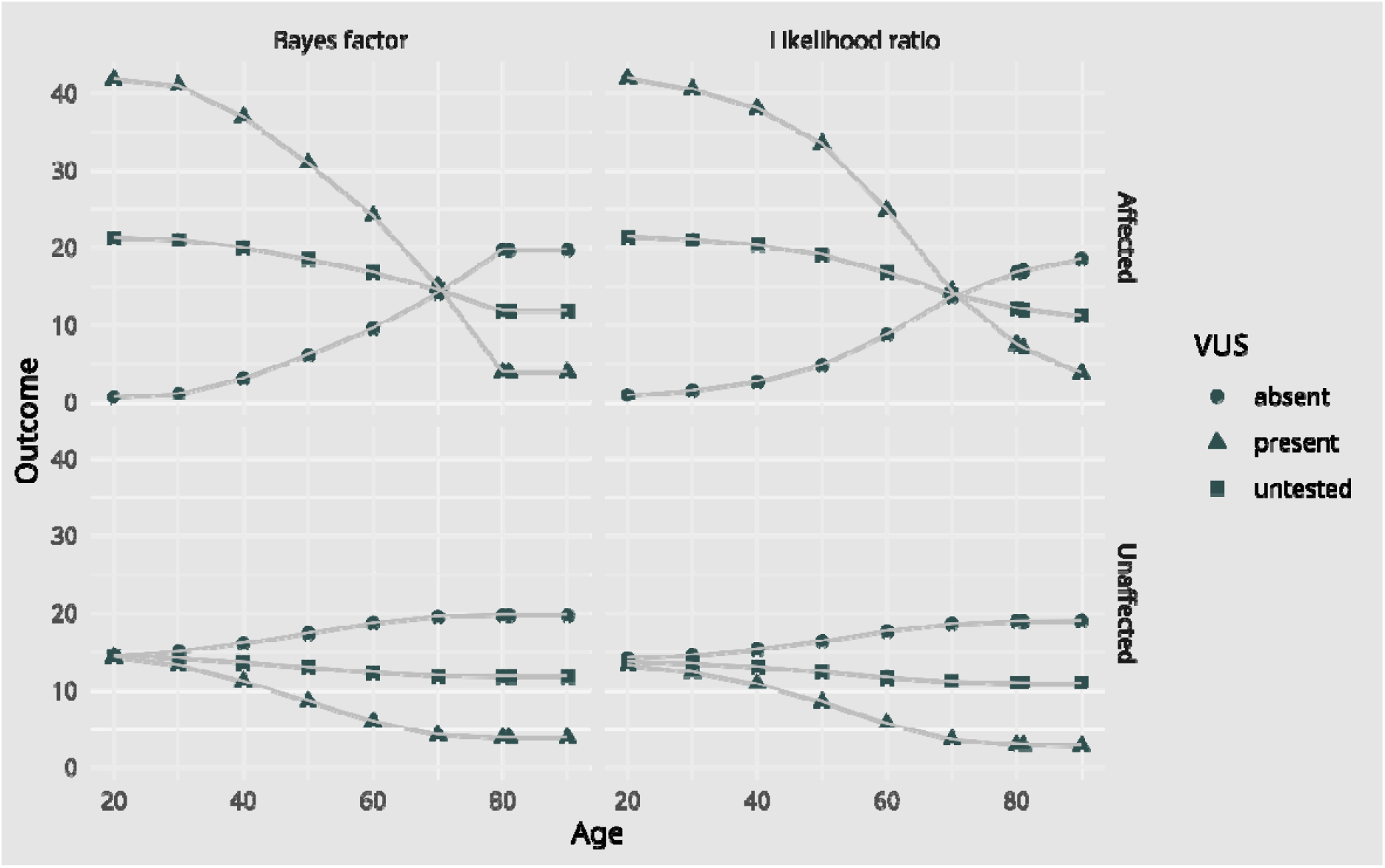
Bayes factor by COOL (5) and LR by CAL-Leiden for person ? in figure 1, at different ages given different affection status and genotype. The default data in COOL and the United Kingdom population penetrance data in CAL-Leiden are used.

Table 1.

**Table 1.**
| Affection status and genotype of person “?” | Rañola et al. (4) | Mohammadi et al. (3) | CAL-Leiden | COOL (5) |
| --- | --- | --- | --- | --- |
|  | analyze.myvariant.org/, as published by Belman et al. | In-house program | https://bioexp.net/cosegregation/ | fengbj-laboratory.org as published by Belman et al. |
| Affected, Variant present | 36.9 | 15.13 | 6.58 | 3.85 |
| Unaffected, Variant present | 9.06 | 6.12 | 2.85 | 3.85 |
| Affected, Variant absent | 17.3 | 24.35 | 15.19 | 19.7 |
| **Unaffected,**<br>**Variant absent** | 42.5 | 30.88 | 16.86 | 19.7 |
| **Affected, Untested** | 27.1 | 19.74 | 10.88 | 11.8 |

Table 1 is based on Belman’s Table S3 (5), in which they compared the performance of their model (COOL) with analyze.myvariant.org from Rañola et al. (4), finding large discrepancies between the two. When we run the original Mohammadi model (3), along with the updated model (CAL-Leiden) and compare them to COOL (5) the results become much more similar. This similarity indicates that the primary differences with the original Mohammadi model (3) are due to outdated penetrance data with higher lifetime risks for carriers, rather than a fundamentally different penetrance model.

Table 1 shows the LRs when affection status and genotype of person ? in pedigree from figure 1 varies. It shows LRs obtained from analyze my variant (as published in Belman et al. (5)), Mohammadi’s co-segregation likelihood model (our in-house model), CAL-Leiden (this paper), and the results from COOL, as published in Belman et al. (5).

When considering the individual as affected or unaffected, along with the presence or absence of the variant or unknown variant status across a grid of ages from 20 to 85, both COOL (5) and CAL-Leiden follow a comparable curve (Figure 2). The slight differences between CAL-Leiden and COOL can be attributed to variations in the underlying population data and differences in the age-specific incidence rates modeling. CAL-Leiden uses a normal-based smoothing approach, while COOL directly employs raw, stepwise constant age-specific age rates. This difference in the modeling of age-specific rates explains the slight divergence between the two models beyond the age of 80. In COOL (5) when the affection status is altered beyond the age of 80, the LR remains numerically equal. This is because COOL enforces a fixed relative risk of cancer of 1 beyond 80 years of age. In contrast, CAL-Leiden, without this constraint due to its smoothing approach, results in closely aligned but decreasing LRs beyond 80.

While the differences in modeling of age-specific incidence rates generally have a small impact on BC, they might significantly affect OC. This is due to important fluctuations in the raw data caused by the lack of events at young ages. Supplementary material figure 1 illustrates the effect of smooth versus stepwise penetrance for all three genes in using a simple pedigree in which OC is present. In the case of *PALB2*, the LR in COOL (5) exhibits significant fluctuations, indicating that the likelihood of pathogenicity is lower when OC occurs between ages 40 and 50 (3.662) compared to diagnosis between ages 50 and 60 (4.489). Furthermore, OC between ages 60 and 70 shows an even lower likelihood (2.266), while diagnosis between ages 70 and 80 exhibits a higher likelihood (2.934).

Additionally, we studied the performance of CAL-Leiden in three pedigrees with varying sizes and structures for *BRCA1, BRCA2* and *PALB2* (supplementary material figure 2). The results demonstrate that the CAL-Leiden model can handle different sizes of pedigrees. Furthermore, the results obtained from CAL-Leiden and COOL (5) exhibit again a high degree of comparability in these pedigrees.

Finally, in supplementary material figure 3, we show how different and multiple within-person cancer types impact the LR in CAL-Leiden within a fixed pedigree configuration for *BRCA1, BRCA2* and *PALB2* genes.

## DISCUSSION

We revised the Mohammadi co-segregation likelihood model (3) by incorporating updated population incidences and relative risks, adding different cancer types other than BC and OC, including the analysis for *PALB2* and updating the modeling for CBC. Furthermore, we have developed a website to facilitate the assessment of co-segregation LR using the penetrance based on the cancer incidence for Dutch or UK population. Regarding our website we have prioritized designing an interface that is exceptionally user-friendly. Upon uploading the pedigree in a CanRisk table format, it is displayed on the screen for visual inspection and confirmation, eliminating the need for manual input. Population incidence for both the UK and NL is also available in dropdown menus, so users don’t need to provide these incidences. After applying the calculation button, the corresponding LR is displayed in the same interface and results can be downloaded as a PDF for personal records. To address patient privacy concerns, especially regarding cross-border data exchange, the model is available upon request. This allows users to process pedigrees within their own institute without uploading data to external servers.

Belman et al. (5) implied that the discrepancies in results between COOL and Mohammadi (3) and Rañola’s (4) model stem from the use of a ‘cumulative risk model’. However, it should be noted that a model based on cumulative risk has never been described by Mohammadi et al. (3). As explained above, both the Mohammadi model (3) and the Rañola adaptation of it (4) rely on a survival approach, similar to the approach used by COOL (5). As shown in the results section, CAL-Leiden and COOL (5) show overall comparable results. The observed slight differences can presumably be attributed to differences in population incidences, relative risks, and differences in the age-specific incidence rates modeling. In CAL-Leiden, the raw data, based on age intervals with constant incidence rates, is smoothed using the best-fitting (least squares) normal approximation. This leads to smooth estimates of the age-specific age rates, reducing spurious fluctuations in the data caused by combining age-dependent relative risks and underlying population incidence rates of non-coinciding intervals. On the other hand, COOL (5) uses the original raw age-specific cancer incidence rates based on intervals. The practical impact on LR of this different approach is small.

In some cases, a person in the pedigree is diagnosed with multiple gene-related cancers, such as BC and OC or CBC. The original model of Mohammadi (3) and CAL-Leiden also considered the occurrence of CBC. In contrast to the Mohammadi’s approach based on independence, in the CAL-Leiden, we apply a more advanced approach using newly available data on the relative risks of CBC given the age and the elapsed time from the first primary BC diagnosis (7).

Since there is limited published data on the risk for individuals with a previous cancer diagnosis across most combinations of cancers, there are essentially three options: consider them independently, discard all information after the first diagnosis or discard all information after the first diagnosis except the CBC after the first BC. CAL-Leiden allows users to consider these options in a user-friendly way, without modifying the input data. If the “Complete” option is chosen in the website tool, all diagnoses of gene-related cancers within a person are considered, treating them as independent processes, except CBC which is considered as explained above. One might prefer, however, to end the follow-up at the first cancer diagnosis. This alternative approach is also available in CAL-Leiden: “First diagnosis”, allows users to choose this option, without modifying the input data. This approach is consistent with the methodology used in COOL (5) and aligns with the current guidelines of the Variant Curation Expert Panel (VCEP) (17, 18). However, such an approach has the downside of discarding potentially relevant information with respect to the pathogenicity of the variant under investigation, leading to underestimation of the variant’s pathogenicity likelihood. For example, when a VUS is identified in a woman with both BC and OC diagnosed at a young age, it is more likely that the variant is pathogenic compared to when she has had only BC. For those who wish to end follow-up after the first cancer diagnosis but want to include CBC after the first BC, the “First diagnosis+CBC” option is available.

The inclusion of prostate cancer in the co-segregation calculations is a subject of ongoing debate. Previous research has firmly linked pathogenic *BRCA2* variants to an increased risk of prostate cancer (11, 19). The European Randomized Study of Screening for Prostate Cancer (ERSPC) demonstrated reduced mortality with PSA-based screening (20, 21), prompting its adoption in certain countries. This raises questions about whether prostate cancer, particularly when detected through screening, introduces bias into risk assessments. Preferably we would like to exclude certain low risk prostate tumors which are diagnosed based on PSA-screening. However, we cannot be sure that the user knows the details of each prostate tumor and can apply it in preparing the data file for calculation. Therefore, we have chosen to not include prostate cancer in the calculations.

Quantitative co-segregation analysis methods are well established for their use in interpretation of causality of *BRCA1, BRCA2* and *PALB2* variants in ACMG/AMP guidelines. The LRs can be converted to different PP1 (supporting strength for pathogenicity) and BS4 (lack of segregation in affected members of a family) strength categories, following the LR thresholds described in Tavtigian et al. (1). If there are independent families with the same variant, the LRs can be multiplied. Differences in LR between different co-segregation models could potentially bear significance when the LR approaches the ACMG threshold for a specific strength criterion. However, it’s worth noting that in many instances, the variances in LR are minimal, rendering this an often inconsequential issue. Given the substantial similarities between the tools, the choice of which model to employ may be influenced by user familiarity, specific (safety) rules and regulations, or other nuanced considerations. Additionally, it is important to recognize that none of the existing models for VUS classification, relying on co-segregation data, currently incorporate measures of uncertainty into their outcomes. This issue becomes particularly significant when the LR is derived from a limited number of families. To address this challenge, it is important to enhance the confidence in the classification by combining the results from unrelated families carrying the same VUS. International collaborations like the ENIGMA consortium facilitate data aggregation, ultimately leading to a more reliable reclassification of VUS. This, in turn, holds utmost significance for patient care and treatment decisions.

## DATA AVAILABILITY STATEMENT

For pedigree outputs in the main manuscript or supplementary material, please contact the authors.

## CODE AVAILABILITY

For the code used in the model, please contact the authors.

## ACKNOWLEDGMENTS

We thank Prof. A. Antoniou (Cancer Research UK, Cambridge, UK) for approval of using the CanRisk data format in CAL-Leiden model.

## AUTHOR CONTRIBUTION STATEMENT

Conceptualization: M.P.G.V., S. M., M.R.G.

Formal analysis: S.M., R.M., M.R.G.

Funding acquisition: M.P.G.V.

Investigation: S.M., M.R.G., M.E.B.

Methodology: M.R.G., S.M., M.P.G.V.

Software: R.M.

Writing-review & editing: S.M., R.M., M.E.B, M.P.G.V., M.R.G.

## ETHICAL APPROVAL

Not applicable. Simulated pedigrees are used.

## REFERENCES

1. Tavtigian SV, Greenblatt MS, Harrison SM, Nussbaum RL, Prabhu SA, Boucher KM, et al. Modeling the ACMG/AMP variant classification guidelines as a Bayesian classification framework. Genet Med. 2018;20(9):1054–60.

2. Thompson D, Easton DF, Goldgar DE. A full-likelihood method for the evaluation of causality of sequence variants from family data. Am J Hum Genet. 2003;73(3):652–5.

3. Mohammadi L, Vreeswijk MP, Oldenburg R, van den Ouweland A, Oosterwijk JC, van der Hout AH, et al. A simple method for co-segregation analysis to evaluate the pathogenicity of unclassified variants; BRCA1 and BRCA2 as an example. BMC Cancer. 2009;9:211.

4. Rañola JMO, Liu Q, Rosenthal EA, Shirts BH. A comparison of cosegregation analysis methods for the clinical setting. Fam Cancer. 2018;17(2):295–302.

5. Belman S, Parsons MT, Spurdle AB, Goldgar DE, Feng BJ. Considerations in assessing germline variant pathogenicity using cosegregation analysis. Genet Med. 2020;22(12):2052–9.

6. Jonker MA, Jacobi CE, Hoogendoorn WE, Nagelkerke NJ, de Bock GH, van Houwelingen JC. Modeling familial clustered breast cancer using published data. Cancer Epidemiol Biomarkers Prev. 2003;12(12):1479–85.

7. Kuchenbaecker KB, Hopper JL, Barnes DR, Phillips KA, Mooij TM, Roos-Blom MJ, et al. Risks of Breast, Ovarian, and Contralateral Breast Cancer for BRCA1 and BRCA2 Mutation Carriers. Jama. 2017;317(23):2402–16.

8. Mocci E, Milne RL, Méndez-Villamil EY, Hopper JL, John EM, Andrulis IL, et al. Risk of pancreatic cancer in breast cancer families from the breast cancer family registry. Cancer Epidemiol Biomarkers Prev. 2013;22(5):803–11.

9. Antoniou AC, Cunningham AP, Peto J, Evans DG, Lalloo F, Narod SA, et al. The BOADICEA model of genetic susceptibility to breast and ovarian cancers: updates and extensions. Br J Cancer. 2008;98(8):1457–66.

10. Thompson D, Easton DF. Cancer Incidence in BRCA1 mutation carriers. J Natl Cancer Inst. 2002;94(18):1358–65.

11. Cancer risks in BRCA2 mutation carriers. J Natl Cancer Inst. 1999;91(15):1310–6.

12. Yang X, Leslie G, Doroszuk A, Schneider S, Allen J, Decker B, et al. Cancer Risks Associated With Germline PALB2 Pathogenic Variants: An International Study of 524 Families. J Clin Oncol. 2020;38(7):674–85.

13. IKNL. NKR Cijfers, Incidentie, prevalentie, sterfte]. https://nkr-cijfers.iknl.nl/#/viewer.

14. Cancer research UK. Health professional, Data and Statistics, Cancer Statistics, Statistics by cancer type]. https://www.cancerresearchuk.org.

15. Peto J, Mack TM. High constant incidence in twins and other relatives of women with breast cancer. Nat Genet. 2000;26(4):411–4.

16. CanRisk. https://www.canrisk.org/.

17. ENIGMA BRCA1 and BRCA2 Variant Curation Expert Panel. https://clinicalgenome.org/affiliation/50087.

18. Hereditary Breast, Ovarian and Pancreatic Cancer Variant Curation Expert Panel. https://clinicalgenome.org/affiliation/50039.

19. Kote-Jarai Z, Leongamornlert D, Saunders E, Tymrakiewicz M, Castro E, Mahmud N, et al. BRCA2 is a moderate penetrance gene contributing to young-onset prostate cancer: implications for genetic testing in prostate cancer patients. British Journal of Cancer. 2011;105(8):1230–4.

20. Kohestani K, Chilov M, Carlsson SV. Prostate cancer screening-when to start and how to screen? Transl Androl Urol. 2018;7(1):34–45.

21. Page EC, Bancroft EK, Brook MN, Assel M, Hassan Al Battat M, Thomas S, et al. Interim Results from the IMPACT Study: Evidence for Prostate-specific Antigen Screening in BRCA2 Mutation Carriers. Eur Urol. 2019;76(6):831–42.

