## Supplementary material for "A Comprehensive and Accessible Model for Co-Segregation Analysis in *BRCA1, BRCA2*, and *PALB2* Variant Classification"

**Content:**

**Appendix A:** Likelihood ratio derivation from CAL-Leiden.

**Appendix B:** Phenotypic likelihood contribution of contralateral breast cancer.

**Appendix C:** Comparison of different phenotypic likelihood formulations.

**Supplementary table 1:** Updated input parameters using population incidence rates in The Netherlands and The United Kingdom.

**Supplementary table 2:** Relative risks.

**Supplementary figure 1:** The effect of smoothing in CAL-Leiden versus stepwise penetrance as used in COOL (1).

**Supplementary figure 2:** The effect of different sizes and complexities and different diagnosis on likelihood ratios in CAL-Leiden and COOL (1).

**Supplementary figure 3:** The effect different diagnosis on likelihood ratio in CAL-Leiden.

**Appendix A. Likelihood ratio derivation from CAL-Leiden**

The likelihood ratio (LR) for pathogenicity of a variant based on segregation within families is defined as follows:

$LR=\frac{P_{d}(G_{o}|{Ph}_{f},G_{p}=1)}{P_{n}(G_{o}|{Ph}_{f},G_{p}=1)}$ ,	(S1)

where *Ph_f_* stands for the phenotypes in family *f*, *G_O_* stands for the observed profile of genotypes of the VUS in family *f,* and *G_p_* is the genotype of the proband, who is a carrier per definition. Note that we assume that the observation of cancer phenotypes in the family is complete but that there is missing genotype information. In general, subscript “*f”* refers to all family members, and can be decomposed into two disjoint sets of observed genotypes indexed by subscript “*o*” and a set of unobserved genotypes, indexed by subscript “*u*”.

Applying Bayes’ rule and total probability law, and denoting by $G_{O}^{'}$ all possible genotype configurations based on the observed data, the numerator of (S1) can be rewritten as:

$P_{d}\left( G_{o} | {Ph}_{f},G_{p}=1 \right)\boldsymbol{=}\frac{P_{d}\boldsymbol{(}{Ph}_{f}\boldsymbol{|}G_{o}\boldsymbol{,}G_{p}=1\boldsymbol{)}P\boldsymbol{(}G_{o}|G_{p}=1\boldsymbol{)}}{\sum_{\boldsymbol{G}_{\boldsymbol{o}}^{\boldsymbol{'}}} P_{d}\boldsymbol{(}{Ph}_{f}\boldsymbol{|}G_{o}^{'}\boldsymbol{,}G_{p}=1\boldsymbol{)}P\boldsymbol{(}G_{o}^{'}|G_{p}=1\boldsymbol{)}}$	(S2)

Now, considering the untested individuals (missing genotype), we can rewrite (S2) as follows:

$P_{d}\left( G_{o} | {Ph}_{f},G_{p}=1 \right)\boldsymbol{=}\frac{\sum_{\boldsymbol{G}_{\boldsymbol{u}}} \left\{ P_{d}\left( {Ph}_{f} | G_{o}\boldsymbol{,}{G_{u},G}_{p}=1 \right)P\left( G_{o},G_{u} | G_{p}=1 \right) \right\}}{\sum_{\boldsymbol{G}_{\boldsymbol{f}}} P_{d}\boldsymbol{(}{Ph}_{f}\boldsymbol{|}G_{f}\boldsymbol{)}P\boldsymbol{(}G_{f}|G_{p}=1\boldsymbol{)}}$	(S3)

On the other hand, under neutrality of the variant, genotype and phenotypes are independent and hence $P_{n}\left( G_{o} | {Ph}_{f},G_{p}=1 \right)=P(G_{0}|G_{p}=1)$.

**Appendix B. Phenotypic likelihood contribution of contralateral breast cancer**

**Phenotypical likelihood contribution of contralateral breast cancer in Mohammadi et al. (10)**

As explained in the main text, the original model in Mohammadi et al. (2009) allowed for the consideration of all primary cancer diagnoses of gene-specific relevant cancer types within a person, assuming their independence. When a person experienced both ovarian cancer (OC) and breast cancer (BC), this was accounted for in expression (2) by multiplying their density functions evaluated at their respective ages at onset. If one or both cancers were not experienced, this was incorporated into the likelihood calculation through the cancer-specific survival function at the age of last follow-up.

The scenario of contralateral breast cancer (CBC), involving two primary breast tumors—one in each breast—requires clarification. Let *t_bci_* denote the age at onset of the primary breast cancer in one breast and denote by *t_cbci_* (*t_cbci_> t_bci_*) denotes the age at onset of breast cancer in the contralateral breast.

When bilateral breast cancer occurs, the available, age-specific, observed breast cancer incidence cannot be directly attributed to each primary breast tumor individually; instead, it refers to their combined effect, specifically to the one that occurred first. Hence, the available cumulative distribution functions (under variant pathogenicity and neutrality assumptions) from the state-of-the-art literature refer to the random variable $\tilde{T}=min(T_{right},T_{left})$, where $T_{right}$ and $T_{left}$ refer, respectively, to the age at breast cancer onset in the right breast and the age at breast cancer onset in the left breast, assumed to be independent and identically distributed. Consequently, based on general results in the theory of random variables, the cumulative distribution function of breast cancer at age *t* in a particular breast can be written as $F_{bc}^{'}\left( t \right)=1-\sqrt{1-F_{bc}(t)}$. As a result, the likelihood for bilateral breast cancers with onset ages *t_bci_* and *t_cbci_* is given by the product of the derivatives of $F_{bc}^{'}\left( t_{bci} \right)$ and $F_{bc}^{'}\left( t_{cbci} \right)$. It is important to note that when contralateral breast cancer was not observed but a primary breast cancer was observed, the probability of not having observed contralateral breast cancer was not considered. In other words, contralateral breast cancer only affected the phenotypic likelihood when observed.

**Phenotypical likelihood contribution of contralateral breast cancer in CAL-Leiden**

Kuchenbaker et al. (3) includes incidence rates of contralateral breast cancer conditional to the age of the primary breast cancer for *BRCA1* and *BRCA2* carriers. This allows us to go beyond the independence assumption and consider contralateral breast cancer in the phenotypical likelihood of CAL-Leiden in a more realistic way.

Specifically, using Bayes’ theorem, we can rewrite the joint density function of *t_cbi_* and *t_cbci_* as follows:

$f_{bc}\left( t_{bci},t_{cbci} \right)=f_{bc}\left( t_{bci} \right)f\left( t_{cbci} | t_{bci} \right)$	(S7)

The term $f\left( t_{cbci} | t_{bci} \right)$ can be rewritten as $f\left( t_{cbci}-t_{bci} | t_{bci} \right)$ in line to how it is presented in publications. Since the data on the incidence of contralateral breast cancer stratified by values of $t_{bci}$ presents high uncertainty in (3), we simplify $f\left( t_{cbci}-t_{bci} | t_{bci} \right)$ and consider $f(t_{cbci}-t_{bci})$ instead.

We applied a smoothing approach for the term $f(t_{cbci}-t_{bci})$ based on an exponential, constant-hazards distribution, given that corresponding raw incidence rates do not clearly vary with age. Like the normal approximations used for the primary gene-related cancers considered, we used least squares to determine the best-fitting exponential distribution with parameters r (lifetime risk) and $\lambda$ (rate). This resulted in parameters r= 0.72 and $\lambda=0.01$ for non-carriers, r=0.66 and $\lambda=0.04$ for *BRCA1* carriers and r=0.52 and $\lambda=0.03$ for *BRCA2* carriers.

Additionally, contralateral breast cancer is treated in CAL-Leiden as any other cancer in the sense that its non-occurrence after a primary breast cancer diagnosis is also taken into account. This is done by considering the probability of not having experienced an additional contralateral breast cancer by the age of end of follow-up through the exponential survival function $S\left( t_{cbci}-t_{bci} \right)$ based on the estimated parameters.

Even though CAL-Leiden no longer relies on the independence assumption for contralateral breast cancer, incorporating it still requires extending the follow-up period and hence considering other non-observed gene-related cancers beyond the first breast cancer diagnosis through the survival function, which does assume independence. While the impact of this is minor, we also provide users the option to include or exclude contralateral breast cancer in the likelihood ratio calculation.

**Appendix C. Comparison of different phenotypic likelihood formulations.**

**Phenotypical likelihood from CAL-Leiden and COOL (9) are equivalent:**

CAL-Leiden is constructed upon a general model that considers age at onset of primary cancers of various types related to the specific genes under consideration (*BRCA1*, *BRCA2*, and *PALB2*), assuming their independence. Given this assumption is arguable, users might prefer to conclude the follow-up at the first cancer diagnosis. CAL-Leiden accommodates this preference through a menu option, which automatically excludes subsequent ages at onset of other cancers from the dataset, eliminating the need for manual data modification. As a result, the age at onset of the remaining relevant cancers is right censored at the age of the first cancer diagnosis. Subsequently, we demonstrate that if follow-up is stopped at the first cancer diagnosis, the underlying phenotypical likelihoods of CAL-Leiden and COOL are equivalent.

Denote by $\tilde{t}$ the earliest occurrence among the cancers of interest and $\tilde{d}=\left\{ 1,\ldots,K \right\}$ an index variable that specifies which event happened first. For example, $\tilde{d}=1$means that breast cancer was the first diagnosed cancer, $\tilde{d}=2$indicates that the firstly diagnosed relevant cancer was ovarian, and so forth. If the individuals did not experience any cancer by the end of follow-up $\tilde{d}=0$.

The CAL-Leiden phenotypical likelihood can be rewritten as:

$P\left( {Ph}_{i} \right)=\left\{ \begin{aligned} f_{k}\left( \tilde{t} \right)\times\prod_{j=1|j\neq k}^{K} (1-F_{j}(\tilde{t})) if \tilde{d}=k \\ \prod_{j=1|j\neq k}^{K} (1-F_{j}(\tilde{t}))if \tilde{d}=0 \end{aligned} \right.$	(S4)

Where $F_{j}(\tilde{t})$ is the cumulative distribution function of cancer type *j* evaluated at age $\tilde{t}$ and $f_{j}\left( \tilde{t} \right)$ its derivative, the density function.

Now, let's turn to the COOL phenotypical likelihood formulation and demonstrate that it is equivalent to expression (S4).

Following Belman et al. (2020), the COOL phenotypical likelihood can be written as:

$P\left( {Ph}_{i} \right)=\left\{ \begin{aligned} h_{k}\left( \tilde{t} \right)\times e^{-\sum_{j=1}^{K} \sum_{t=0}^{\tilde{t}} h_{j}\left( t \right)} if \tilde{d}=k \\ e^{-\sum_{j=1}^{K} \sum_{t=0}^{\tilde{t}} h_{j}\left( t \right)} if \tilde{d}=0 \end{aligned} \right.$	(S5)

Where $h_{j}\left( \tilde{t} \right)$ is the hazard function of cancer type *j* evaluated at age $\tilde{t}$, indicating the probability of occurrence of cancer type *j* at that specific age for individuals who are free of cancer type *j* up to that age. In COOL, the hazard functions $h_{j}$ are approximated using interval-based incidence rates (9).

Expressions (S4) and (S5) are equivalent given the following standard survival analysis results which relate cumulative distribution, survival and cumulative hazards functions:

$S_{j}\left( \tilde{t} \right)=1-F_{j}\left( \tilde{t} \right)$ is the survival function of the time (in age scale) of diagnosis of cancer type *j* evaluated at time $\tilde{t}$ and can be rewritten as follows: $S_{j}\left( \tilde{t} \right)=e^{-\sum_{t=0}^{\tilde{t}} h_{j}(t)}$ given the relation between survival and cumulative hazard functions. On the other hand, the density function of time (in age scale) to cancer type j evaluated at time $\tilde{t}$ can be rewritten as $f_{j}\left( \tilde{t} \right)=h_{j}(\tilde{t})S_{j}(\tilde{t})$. These two equivalences allow to rewrite expression (S4) as (S5) and vice versa.

The only difference between the phenotypical likelihoods of CAL-Leiden and COOL is about the choice for the cancer-specific hazard functions $h_{j}$. As mentioned, COOL directly incorporates 5-year interval-based incidence rates by combining 5-year age-specific population incidence rates and relative risks reported in recent literature. In contrast, CAL-Leiden smooths these incidence rates and utilizes the best normal approximation of the observed rates, optimized using the least squares criterion.

**Phenotypical likelihood implemented in CoSeg R package (2) differs from COOL (1) and CAL-Leiden**

The phenotypical likelihood used in the CoSeg R package is a simplified version of the original phenotypical likelihood proposed by Mohammadi et al. (2009) (10) displayed in equation (2) in the main text. The phenotypical likelihood implemented in CoSeg considers an individual to be affected if they have had either breast or ovarian cancer. The age at onset $\tilde{t}$ is taken as the earliest diagnosis of either breast or ovarian cancer. Furthermore, the formulation exclusively relies on breast cancer penetrance from existing literature, neglecting ovarian cancer penetrance. Expression (S4) is simplified as follows:

$P\left( {Ph}_{i} \right)=\left\{ \begin{aligned} f_{1}\left( \tilde{t} \right) if \tilde{d}=1 or \tilde{d}=2 \\ {1-F}_{1}\left( \tilde{t} \right) if \tilde{d}=0 \end{aligned} \right.$	(S6)

**Supplementary table 1.**

| **Parameters** | | **Female**  **BC** | **Male**  **BC** | **Female**  **OC** | **Female**  **PaC** | **Male**  **PaC** |
| --- | --- | --- | --- | --- | --- | --- |
|  | | **Non carriers** | | | | |
| **μ** | NL | 66.55 | 94.77 | 86.51 | 82.89 | 86.24 |
|  | UK | 70.16 | 98.19 | 89.62 | 101.97 | 95.79 |
| **σ** | NL | 17.99 | 22.15 | 20.13 | 15.02 | 15.75 |
|  | UK | 19.33 | 22.36 | 25.70 | 20.66 | 19.69 |
| **r** | NL | 0.16 | 0 | 0.03 | 0.03 | 0.04 |
|  | UK | 0.18 | 0 | 0.05 | 0.03 | 0.07 |
|  | | **Carriers of a pathogenic *BRCA1* variant** | | | | |
| **μ** | NL | 41.56 | 75.94 | 63.18 | 96.58 | 86.30 |
|  | UK | 45.13 | 75.56 | 60.19 | 114.21 | 92.85 |
| **σ** | NL | 18.32 | 13.97 | 7.07 | 22.66 | 16.89 |
|  | UK | 18.30 | 12.36 | 9.19 | 26.31 | 20.31 |
| **r** | NL | 0.79 | 0.01 | 0.34 | 0.07 | 0.06 |
|  | UK | 0.76 | 0.01 | 0.49 | 0.16 | 0.08 |
|  | | **Carriers of a pathogenic *BRCA2* variant** | | | | |
| **μ** | NL | 47.90 | 74.32 | 62.61 | 78.31 | 79.78 |
|  | UK | 50.39 | 74.35 | 60.45 | 85.03 | 80.36 |
| **σ** | NL | 17.24 | 13.25 | 8.81 | 13.52 | 13.42 |
|  | UK | 17.45 | 11.73 | 9.66 | 15.80 | 14.13 |
| **r** | NL | 0.76 | 0.13 | 0.15 | 0.04 | 0.05 |
|  | UK | 0.75 | 0.10 | 0.22 | 0.06 | 0.05 |
|  | | **Carriers of a pathogenic *PALB2* variant** | | | | |
| **μ** | NL | 56.78 | 80.38 | 92.26 | 74.24 | 132.01 |
|  | UK | 58.40 | 79.60 | 86.61 | 78.78 | 75.63 |
| **σ** | NL | 15.74 | 16.87 | 23.88 | 11.96 | 28.20 |
|  | UK | 16.01 | 14.97 | 26.14 | 13.90 | 5.85 |
| **r** | NL | 0.65 | 0.02 | 0.11 | 0.04 | 1.09 |
|  | UK | 0.65 | 0.02 | 0.13 | 0.04 | 0.04 |

Updated input parameters using population incidence rates in The Netherlands from 2017 and in The United Kingdom from 2015 to 2017.(3, 4)

Values for mean age at diagnosis (μ), standard deviation (σ) and lifetime risk (r) for occurrence of breast cancer (BC), ovarian cancer (OC) and Pancreatic cancer (PaC).

**Supplementary table 2**

| **Gene** | ***BRCA1*** | | | | |
| --- | --- | --- | --- | --- | --- |
|  | **Breast cancer** | | **Ovarian cancer** | **Pancreatic cancer** | |
| **Age** | **Female** | **Male** |  | **Female** | **Male** |
| **0-29** | 73.7 | 8 | 1 | 4.68 | 4.68 |
| **30-39** | 46.2 | 8 | 41.4 | 4.68 | 4.68 |
| **40-49** | 17.2 | 8 | 56.7 | 4.68 | 4.68 |
| **50-59** | 9.7 | 8 | 53.3 | 1.40 | 1.40 |
| **60-64** | 7 | 8 | 53.3 | 1.40 | 1.40 |
| **65-69** | 7 | 8 | 69.1 | 1.40 | 1.40 |
| **70-79** | 4.8 | 8 | 11.8 | 1.40 | 1.40 |
| **Gene** | ***BRCA2*** | | | | |
|  | **Breast cancer** | | **Ovarian cancer** | **Pancreatic cancer** | |
| **Age** | **Female** | **Male** |  | **Female** | **Male** |
| **0-29** | 60.8 | 80 | 1 | 4.77 | 4.77 |
| **30-39** | 20.3 | 80 | 7.3 | 4.77 | 4.77 |
| **40-49** | 16.4 | 80 | 15.9 | 4.77 | 4.77 |
| **50-59** | 11.4 | 80 | 24.5 | 2.03 | 2.03 |
| **60-64** | 6.4 | 80 | 21.5 | 2.03 | 2.03 |
| **65-69** | 6.4 | 80 | 21.5 | 2.03 | 2.03 |
| **70-79** | 6.6 | 80 | 4.4 | 2.03 | 2.03 |
| **Gene** | ***PALB2*** | | | | |
|  | **Breast cancer** | | **Ovarian cancer** | **Pancreatic cancer** | |
| **Age** | **Female** | **Male** |  | **Female** | **Male** |
| **0-29** | 15.6 | 1 | 1 | 1 | 2.67 |
| **30-39** | 10.5 | 6.74 | 2.89 | 1.81 | 3.47 |
| **40-49** | 9.71 | 9.8 | 2.68 | 2.58 | 2.40 |
| **50-59** | 7.45 | 11.7 | 4.34 | 2.27 | 2.94 |
| **60-64** | 6.64 | 7.70 | 1.86 | 2.63 | 1.41 |
| **65-69** | 6.64 | 7.70 | 1.86 | 2.63 | 1.41 |
| **70-79** | 5.9 | 10 | 3.16 | 1.91 | 3.15 |

Relative risks(5-10)

**Supplementary figure 1**

The effect of smoothing in CAL-Leiden versus stepwise penetrance as used in COOL in a simple pedigree (A). Likelihood ratios were calculate based on the age at which ovarian cancer is diagnosed in the niece (marked by the red arrow) of the proband (marked by the black arrow) for a given variant in either *BRCA1*, *BRCA2* or *PALB2* genes. Likelihood ratios are displayed in the table (B) and graphics are presented in section C.

The default data in COOL and the United Kingdom population penetrance data in CAL-Leiden are used.

**A.**

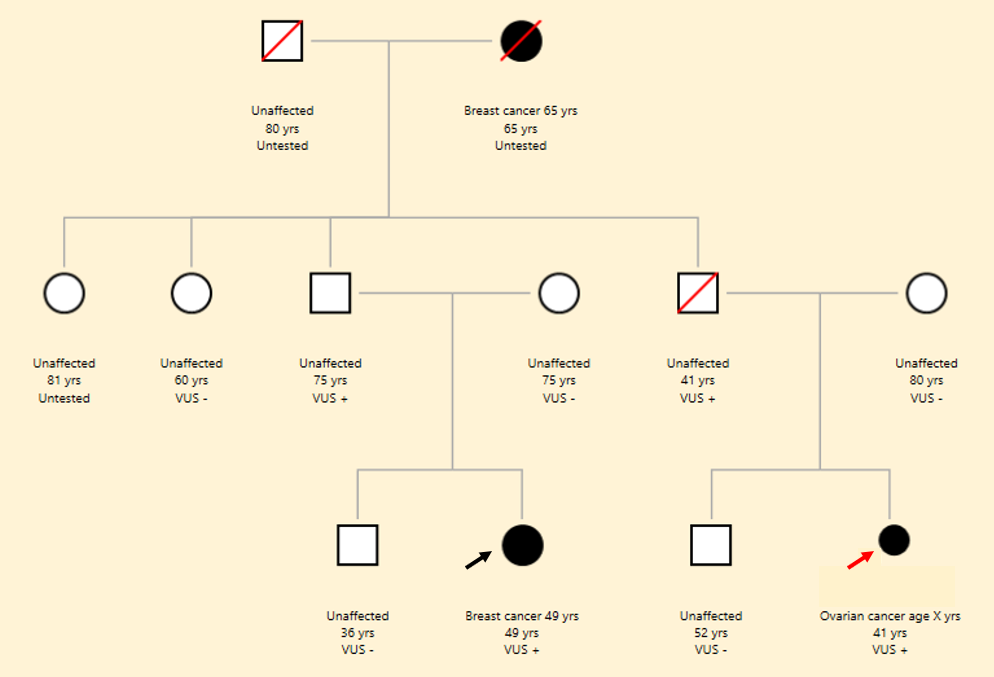

**B.**

|  | ***BRCA1*** | | ***BRCA2*** | | ***PALB2*** | |
| --- | --- | --- | --- | --- | --- | --- |
| **Ovarian cancer at age** | **CAL-Leiden** | **COOL** | **CAL-Leiden** | **COOL** | **CAL-Leiden** | **COOL** |
| **20** | 0.149 | 2.365 | 0.141 | 1.779 | 4.344 | 1.859 |
| **25** | 0.792 | 2.365 | 0.624 | 1.779 | 4.252 | 1.859 |
| **30** | 2.818 | 11.023 | 0.984 | 6.361 | 4.163 | 3.935 |
| **35** | 6.104 | 11.023 | 4.297 | 6.361 | 4.077 | 3.935 |
| **40** | 8.779 | 11.111 | 6.580 | 8.057 | 3.995 | 3.662 |
| **45** | 10.185 | 11.111 | 8.072 | 7.853 | 3.916 | 3.662 |
| **50** | 10.795 | 10.646 | 8.824 | 8.438 | 3.841 | 4.489 |
| **55** | 10.990 | 10.646 | 9.075 | 8.081 | 3.769 | 4.489 |
| **60** | 10.900 | 10.315 | 8.931 | 7.339 | 3.700 | 2.266 |
| **65** | 10.465 | 10.315 | 8.330 | 6.999 | 3.635 | 2.266 |
| **70** | 9.401 | 4.793 | 7.056 | 2.685 | 3.573 | 2.934 |
| **78** | 7.195 | 4.793 | 4.947 | 2.496 | 3.514 | 2.934 |
| **80** | 3.917 | 0.548 | 2.524 | 0.627 | 3.458 | 1.010 |
| **85** | 1.305 | 0.548 | 0.875 | 0.627 | 3.406 | 1.010 |

**C.**

BRCA1

Likelihood ratio

| Ovarian cancer at age (years) | COOL | CAL-Leiden |
| --- | --- | --- |
| 20 | 2.36454 | 0.14914 |
| 25 | 2.36454 | 0.79173 |
| 30 | 11.023 | 2.81758 |
| 35 | 11.023 | 6.1036 |
| 40 | 11.1111 | 8.779 |
| 45 | 11.1111 | 10.185 |
| 50 | 10.6458 | 10.7953 |
| 55 | 10.6458 | 10.99 |
| 60 | 10.3153 | 10.9 |
| 65 | 10.3153 | 10.465 |
| 70 | 4.7931 | 9.4007 |
| 75 | 4.79311 | 7.1952 |
| 80 | 0.54826 | 3.917 |
| 85 | 0.54826 | 1.3051 |

BRCA2

Likelihood ratio

| Ovarian cancer at age (years) | COOL | CAL-Leiden |
| --- | --- | --- |
| 20 | 1.7792 | 0.141 |
| 25 | 1.77923 | 0.624 |
| 30 | 6.36118 | 0.984 |
| 35 | 6.36118 | 4.297 |
| 40 | 8.05725 | 6.58 |
| 45 | 7.8533 | 8.072 |
| 50 | 8.43782 | 8.824 |
| 55 | 8.08064 | 9.075 |
| 60 | 7.3392 | 8.931 |
| 65 | 6.99908 | 8.33 |
| 70 | 2.68464 | 7.056 |
| 75 | 2.49559 | 4.947 |
| 80 | 0.626714 | 2.524 |
| 85 | 0.626711 | 0.875 |

PALB2

Likelihood ratio

| Ovarian cancer at age (years) | COOL | CAL-Leiden |
| --- | --- | --- |
| 20 | 1.85949 | 4.3444 |
| 25 | 1.8595 | 4.25205 |
| 30 | 3.93548 | 4.163 |
| 35 | 3.93548 | 4.07736 |
| 40 | 3.66216 | 3.99514 |
| 45 | 3.66216 | 3.91634 |
| 50 | 4.48901 | 3.84094 |
| 55 | 4.48901 | 3.7689 |
| 60 | 2.26595 | 3.70028 |
| 65 | 2.26595 | 3.6349 |
| 70 | 2.93353 | 3.57295 |
| 75 | 2.93353 | 3.5141 |
| 80 | 1.00955 | 3.458 |
| 85 | 1.00955 | 3.4062 |

**Supplementary figure 2:**

The effect of different sizes and complexities and different diagnosis on likelihood ratios in CAL-Leiden and COOL. Pedigree 1 has 45 family members. Pedigree 2 presents generally the same structure as pedigree 1 but is larger, with 66 family members. These two pedigrees serve as examples of large families. Pedigree 3 is a pedigree with 18 family members and breast and ovarian cancer are present in the family (A). Likelihood ratio results from analysis in CAL-Leiden versus bayes factor from analysis in COOL for *BRCA1*, *BRCA2*, and *PALB2*.

The default data in COOL and the United Kingdom population penetrance data in CAL-Leiden are used.

**A.**

**Pedigree 1, 45 members**

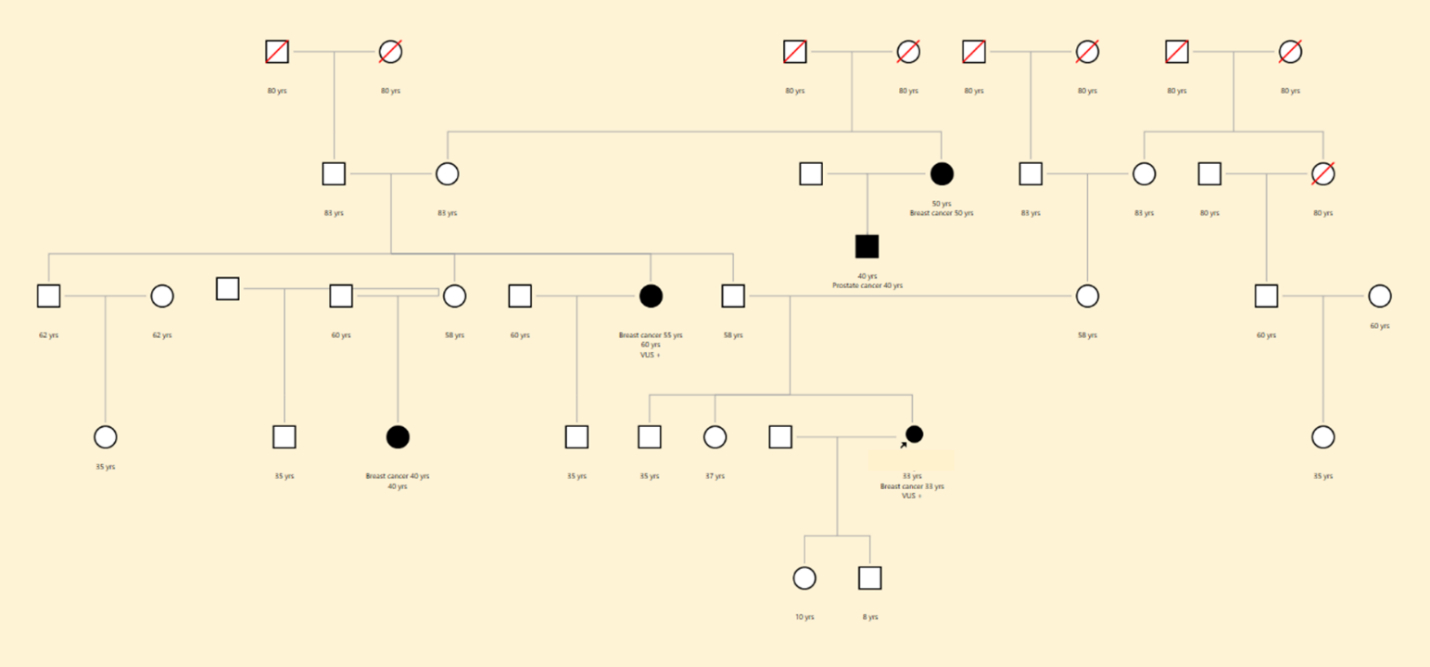

**Pedigree 2, 66 members**

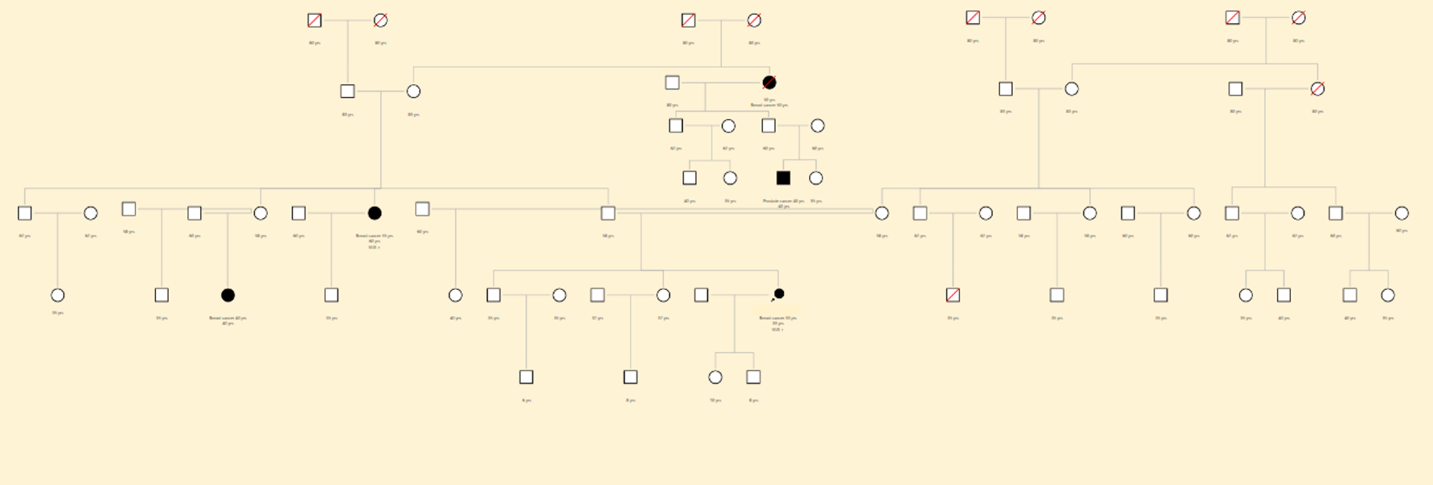

**Pedigree 3, 18 members**

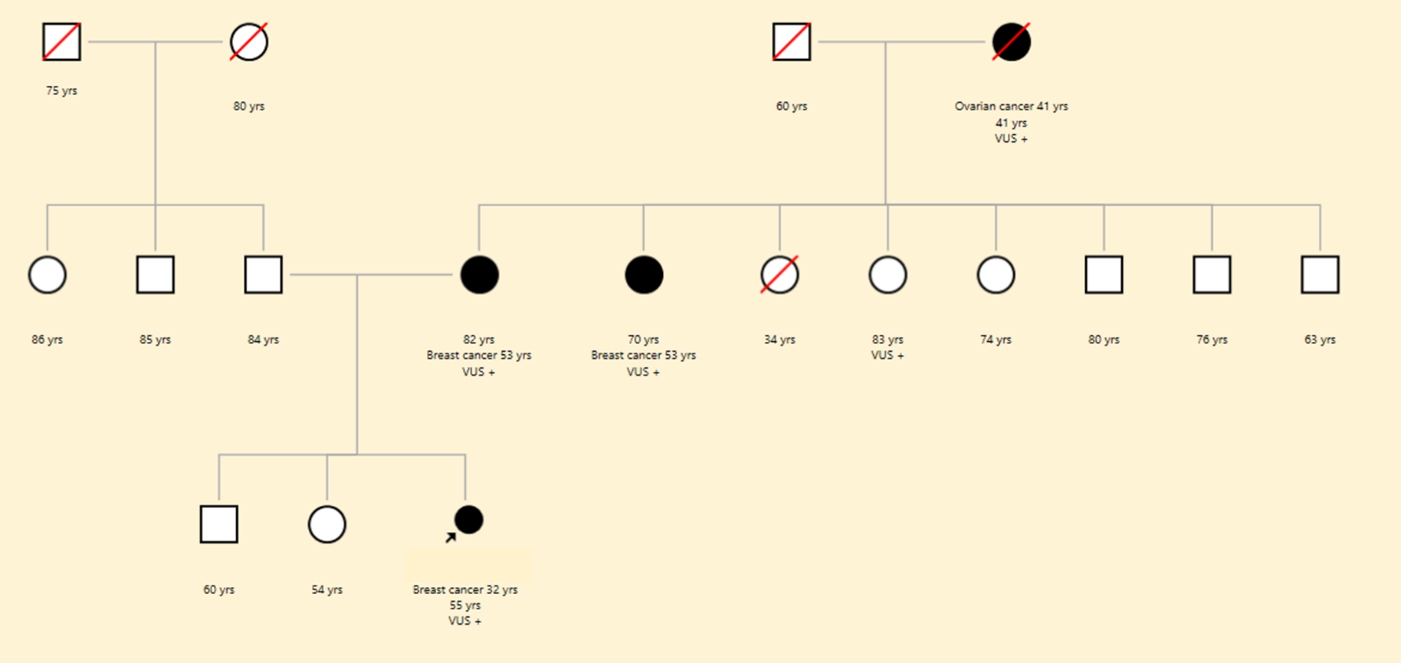

**B.**

|  | **BRCA1** | | **BRCA2** | | **PALB2** | |
| --- | --- | --- | --- | --- | --- | --- |
|  | **CAL-Leiden** | **COOL** | **CAL-Leiden** | **COOL** | **CAL-Leiden** | **COOL** |
| **Pedigree 1** | 2.98 | 3.17 | 3.24 | 3.41 | 3.27 | 3.24 |
| **Pedigree 2** | 3.08 | 3.27 | 3.31 | 3.43 | 3.31 | 3.29 |
| **Pedigree 3** | 1.34 | 2.13 | 2.25 | 3.16 | 3.08 | 3.13 |

**Supplementary figure 3:**

This pedigree is used to demonstrate the effect of different phenotype (combinations) on the likelihood ratio calculated by CAL-L when the multiple cancers option is chosen. In this figure (A) for the person with the red arrow, who is carrier of the VUS, gender and phenotype changes. Genotype remains unchanged. Results (B) vary when the gender and phenotype changes.

The United Kingdom population penetrance data in CAL-Leiden are used.

**A.**

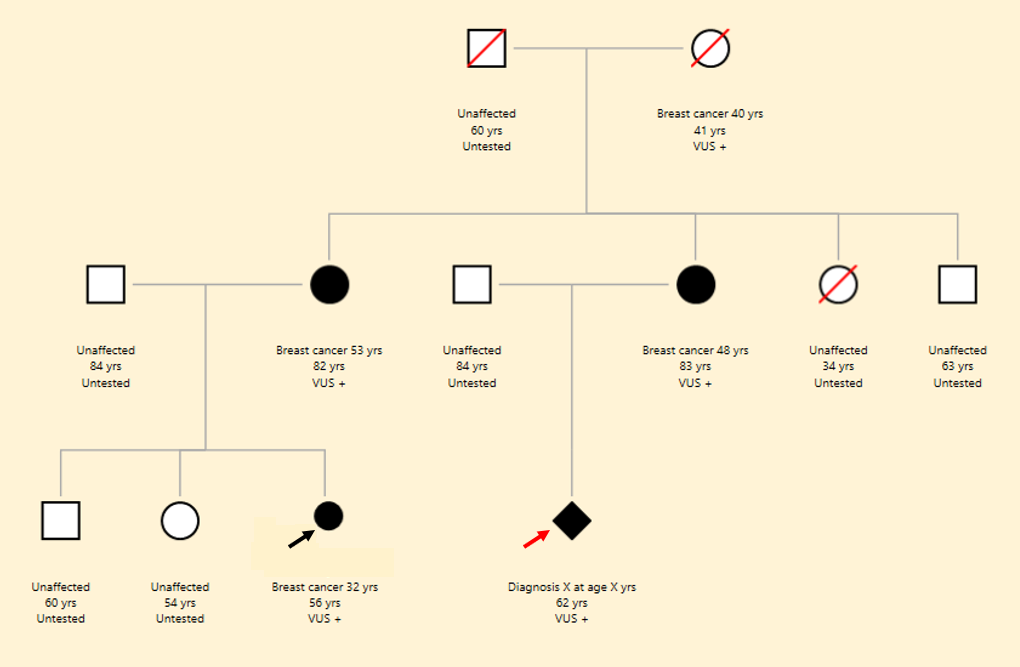

**B.**

| **Age of diagnosis of person with red arrow** | | ***BRCA1*** | ***BRCA2*** | ***PALB2*** |
| --- | --- | --- | --- | --- |
| **Female** | **BC at 45 years** | 11.50 | 12.79 | 12.26 |
|  | **OC at 45 years** | 13.14 | 12.40 | 8.83 |
|  | **PaC at 45 years** | 4.27 | 5.28 | 7.16 |
|  | **CBC at 45 and 62 years** | 13.59 | 13.63 | 14.15 |
|  | **BC and OC**  **respectively at 45 and 62 years** | 14.90 | 14.77 | 13.44 |
|  | **BC and PaC**  **respectively at 45 and 62 years** | 12.54 | 13.86 | 13.38 |
|  | **BC, OC and PaC**  **respectively at 45, 62 and 62 years** | 14.91 | 14.83 | 13.88 |
| **Male** | **BC at 45 years** | 9.65 | 14.20 | 11.71 |
|  | **BC and PaC**  **respectively at 45 and 62 years** | 11.03 | 14.56 | 10.20 |

**References**

1.	Belman S, Parsons MT, Spurdle AB, Goldgar DE, Feng BJ. Considerations in assessing germline variant pathogenicity using cosegregation analysis. Genet Med. 2020;22(12):2052-9.

2.	Rañola JMO, Liu Q, Rosenthal EA, Shirts BH. A comparison of cosegregation analysis methods for the clinical setting. Fam Cancer. 2018;17(2):295-302.

3.	IKNL. NKR Cijfers, Incidentie, prevalentie, sterfte]. <https://nkr-cijfers.iknl.nl/#/viewer>.

4.	Cancer research UK. Health professional, Data and Statistics, Cancer Statistics, Statistics by cancer type]. <https://www.cancerresearchuk.org>.

5.	Kuchenbaecker KB, Hopper JL, Barnes DR, Phillips KA, Mooij TM, Roos-Blom MJ, et al. Risks of Breast, Ovarian, and Contralateral Breast Cancer for BRCA1 and BRCA2 Mutation Carriers. Jama. 2017;317(23):2402-16.

6.	Mocci E, Milne RL, Méndez-Villamil EY, Hopper JL, John EM, Andrulis IL, et al. Risk of pancreatic cancer in breast cancer families from the breast cancer family registry. Cancer Epidemiol Biomarkers Prev. 2013;22(5):803-11.

7.	Antoniou AC, Cunningham AP, Peto J, Evans DG, Lalloo F, Narod SA, et al. The BOADICEA model of genetic susceptibility to breast and ovarian cancers: updates and extensions. Br J Cancer. 2008;98(8):1457-66.

8.	Thompson D, Easton DF. Cancer Incidence in BRCA1 mutation carriers. J Natl Cancer Inst. 2002;94(18):1358-65.

9.	Cancer risks in BRCA2 mutation carriers. J Natl Cancer Inst. 1999;91(15):1310-6.

10.	Yang X, Leslie G, Doroszuk A, Schneider S, Allen J, Decker B, et al. Cancer Risks Associated With Germline PALB2 Pathogenic Variants: An International Study of 524 Families. J Clin Oncol. 2020;38(7):674-85.
